# A New In-Silico Approach for PROTAC Design and Quantitative Rationalization of PROTAC mediated Ternary Complex Formation

**DOI:** 10.1101/2022.07.11.499663

**Authors:** A S Ben Geoffrey, Nagaraj M Kulkarni, Deepak Agrawal, Rajappan Vetrivel, Kishan Gurram

## Abstract

Proteolysis-Targeting Chimeric Molecules (PROTAC) is a rapidly emerging technology for drug target protein degradation and drugging undruggable drug targets. There is a growing literature on in-silico approaches to the complex problem of PROTAC design, specifically the advantages of AI/ in-silico methods in the PROTAC Design Make Test Analyze (DMTA) cycle. Our work presented here aims to contribute to the growing literature of in-silico approaches to PROTAC design by incorporating and demonstrating incremental advancement over previously published methods. We use AI based generative methods for PRTOAC design and Molecular Dynamics to evaluate the stability of the ternary complex formed and ability of the PROTAC to hold the target protein and E3 ligase together stably. To quantify the performance of the PROTAC candidate, we also estimate computationally the PROTAC performance metrics routinely measured by the experimentalists in PROTAC assays. We use highly accurate absolute binding free energy calculations used traditionally in protein-ligand space for the PROTAC system. We calculate (Gibbs free energy change) ΔG for binary complex formation and ternary complex formation mediated by the PROTAC using Free Energy Perturbation - Thermodynamics Integration (FEP-TI) method which is benchmarked in literature with a root mean square error of 0.8 kcal/mol. We calculate ΔG for ternary and binary complexes and estimate whether ΔG for ternary is lower than the ΔG estimated for binary complexes. When the ΔG for ternary is lower than the binary it is inferred that ternary complexation is favoured over binary. Therefore, through these methods we can theoretical estimate ΔG measured by experimentalists in PROTAC assays such as Isothermal titration calorimetry (ITC) and Surface plasmon resonance (SPR) which capture the ΔG for ternary and binary complex formation mediated by the PROTAC. This method will help reduce time as well as costs of the PROTAC DMTA cycle and will accelerate early stage PROTAC drug discovery. As an illustrative application of our in-silico PROTAC design approach, we chose the target Fibroblast growth factor receptor 1(FGFR-1) which is a target approved drug for colorectal cancer. We report the findings and conclude with future research directions.

## Introduction

Proteolysis-Targeting Chimeric Molecules (PROTAC) is a rapidly emerging technology for drug target protein degradation and drugging undruggable drug targets. There is a growing literature on in-silico approaches to the complex problem of PROTAC design, specifically the advantages of AI/ in-silico methods in the PROTAC Design Make Test Analyze (DMTA) cycle.

In-silico approaches to rationalize PROTAC mediated ternary complex formation begin with the work of Drummond et.al. (2019) [1] where they use protein-protein docking and conformation search of PROTAC candidates to rationalize PROTAC mediated ternary complex formation. This we call as the first generation of methods. The use of Molecular Dynamics simulations to access the stability of the PROTAC mediated ternary complex was used by Testa et.al (2020) [2]. Though the MD approach enables us to assess whether the PROTAC candidate can mediate a stable ternary complex formation involving the target protein and E3 ligase, it does not provide a quantifiable measure to rank different PROTAC candidates on their ability to mediate a stable ternary complex formation.

This deficiency has been addressed in the next generation of methods which begin to appear in the 2022 literature. Liao, Junzhuo, et al (2022) [3] and Li, Wenqing, et al (2022) [4] have reported methods that use molecular mechanics combined with the generalized Born and surface area continuum solvation (MMGBSA) and Molecular Mechanics Poisson-Boltzmann Surface Area (MMPBSA) calculated ΔGs to relatively rank PROTAC candidates and achieved a good correlation (R^2^ > 0.9) with ranking based on experimentally measured quantities from PROTAC assays. This we call as the third generation of methods.

While relative ranking based on in silico approach is possible, so far, to our knowledge our work is the first work which reports absolute free energy change of binary and ternary complex formations and thereby computes theoretically the metrics measured by experimentalists in PROTAC assays including Roy et al (2019) [5], Zhenyi and Crews (2022) [6]. The method we used to calculate the absolute free energy change of binding is reported in the literature to have a root mean square error of 0.8 kcal/mol (Chemical Science (2016) [7]) which is the best result so far. Additionally, we also leverage the recent developments of generative AI based linker design for PROTACs in our in-silico workflow (Imrie et al (2020) [8]). To illustrate our in-silico workflow for PROTAC design, we implement our workflow to design a PROTAC for FGFR-1 target with an approved small molecule drug for colorectal cancer.

## Methodology

PROTAC design using our approach was conducted for the target protein, FGFR-1, which is implicated in cancer causing pathways and is overexpressed and validated target for colorectal cancer. We chose Murine Double Minute 2 (MDM2) as the E3 ligase of choice for tagging FGFR-1 for degradation. The support for the use of MDM2 as E3 ligase of choice in colorectal cancer is present in Hines et al (2019) [9].

### Step 1

The first step of the approach is as follows. We dock the known binders of FGFR-1 and MDM2 to their respective targets and perform protein-protein docking of FGFR-1 and MDM2 complexes. We identify protein-protein docking poses with the ligands in physically close proximity that allow them to be connected by a linker.

### Step 2

We use the approach of Imrie et al (2020)[8] of deep generative model for PROTAC linker design and align the generated PROTAC molecule with the two positions of the docked ligands in FGFR-1 --- MDM2 docked pose and identify the PROTAC candidates having the minimum Root Mean Square Deviation (RMSD) of alignment, so that the interactions of the ligands with FGFR-1 and MDM-2 are intact. Our generative model resulted in a total of forty-eight different PROTAC candidates with different linkers.

### Step 3

Next step in the workflow involves scoring/ranking the PROTAC candidates to choose a PROTAC candidate with most interactions with the pocket residues of FGFR-1 and MDM2. To do this, unlike Li, Wenqing, et al. (2022) [4] who used a MMGBSA which is traditionally used for scoring protein-ligand interactions for scoring PROTAC-protein interactions, we instead used a molecular docking-based approach to score/rank the PROTAC candidates. We generate multiple conformers of the PROTAC candidate in the pocket formed at the interface of FGFR1-MDM2 and score the interaction of a given PROTAC conformers with the residues in the pocket using pretrained Deep Learning models which were trained on PDBbind dataset to predict binding affinity between organic molecules and protein residues (Rezaei et al (2020) [10]). The required input configuration of the pocket residues of both the proteins (FGFR1- MDM2) and a single PROTAC conformer to predict the interaction is shown in Fig.1 below.

**Fig. 1.**
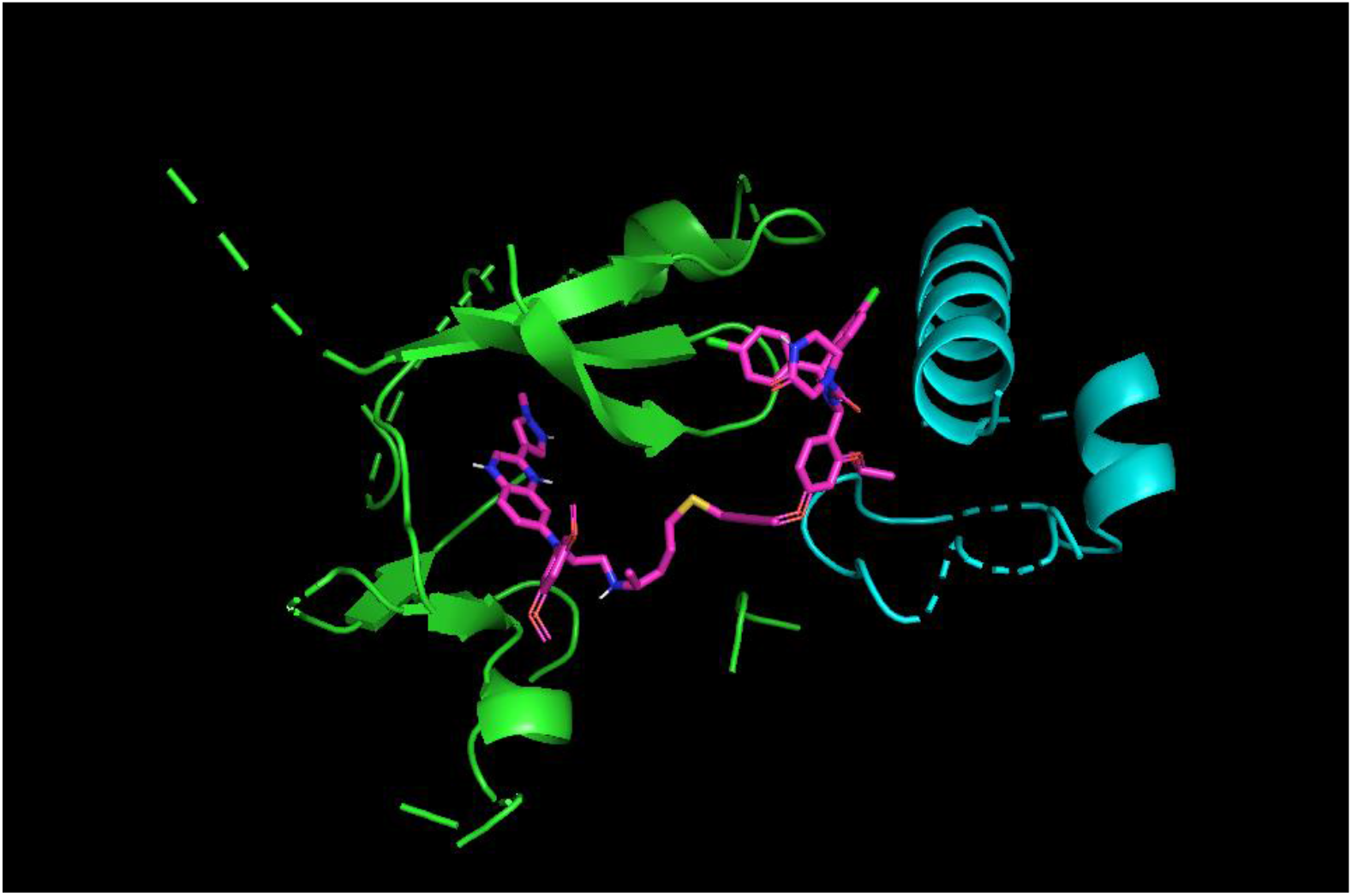
Pocket residues of FGFR1-MDM2 and a PROTAC conformer

### Step 4

After the initial scoring and ranking of PROTAC candidates we select the top-ranking candidates for analysis using more computationally expensive methods in our workflow. We use the top ranking PROTAC candidate to conduct a Molecular Dynamics simulation to access the stability of the formed PROTAC mediated ternary complex. We analyse whether the interactions are stable and intact after a 200 ns long simulation run. Further, while Li, Wenqing, et al (2022) [4] have adopted MMGBPSA which is traditionally used for calculating ΔG associated with protein-ligand binding to calculate ΔG of ternary and binary complex formation and infer whether the stable formation of ternary complex is thermodynamically favoured, we use Free Energy Perturbation – Thermodynamics Integration method which is benchmarked in literature with a root mean square error of 0.8 kcal/mol for ΔG calculations. We estimated the ΔG for the ternary complex formation involving the PROTAC and FGFR-1 --- MDM2 and the ΔG for the binary complexes involving the PROTAC and FGFR-1 followed by PROTAC and MDM2. We estimate whether ΔG for ternary is lower than the binary which would indicate stable ternary complex formation. We do so by decoupling the PROTAC in forty lambda windows as it is conducted in FEP simulations. The entire workflow of the four main steps of our method is captured in the flow diagram given below in Fig.2.

**Fig. 2.**
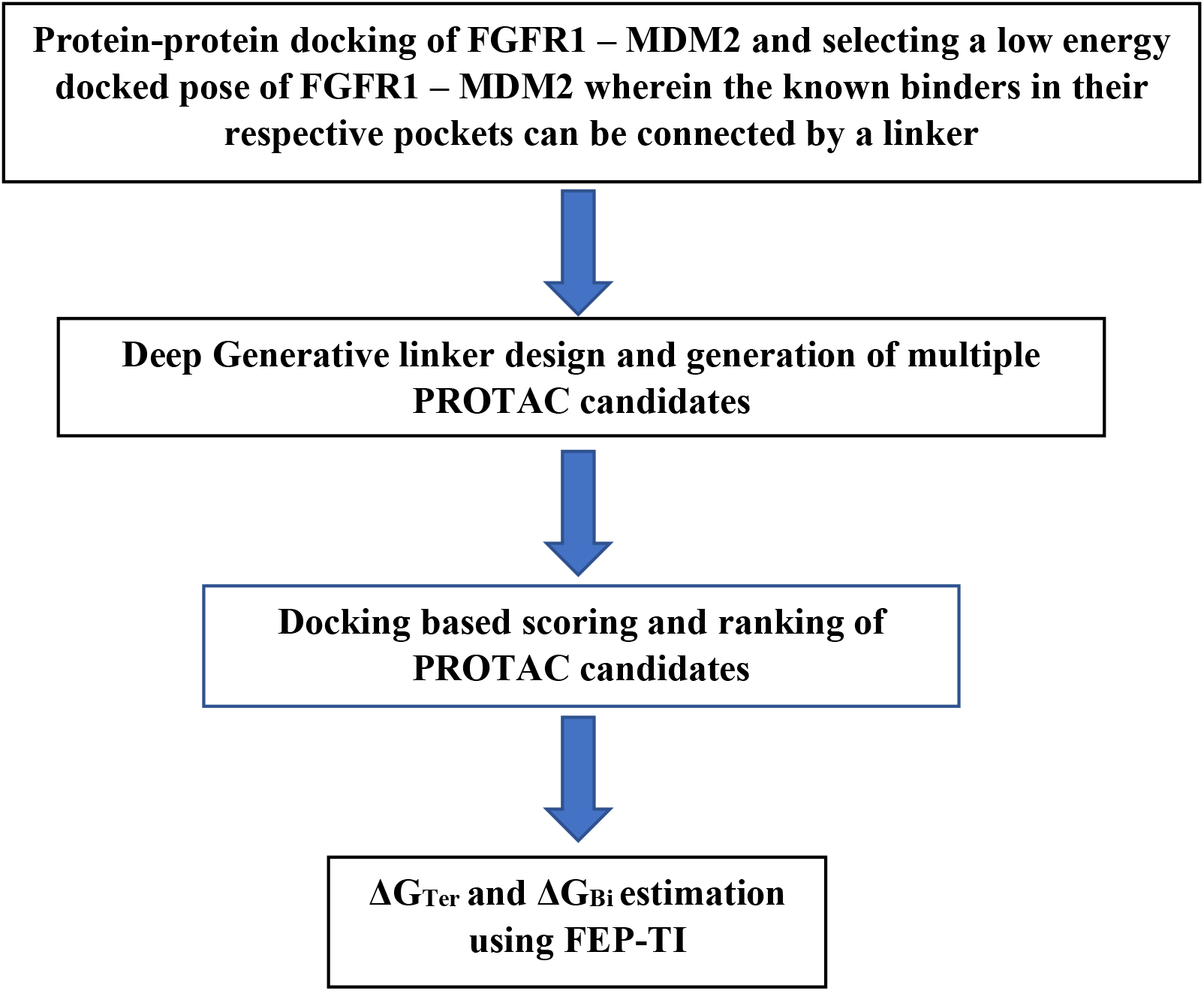
Workflow diagram.

## Results and discussion

Overall, we generated forty-eight different PROTAC candidates for mediating interaction between FGFR1 and MDM2. We scored and ranked them based on our docking-based scoring method. The top seven among the forty-eight different PROTAC candidates designed with different linkers by our AI based linker design approach are listed in Table 1 below. The table captures the structure, SMILES, and the score from our scoring (docking) method.

**Table 1.**
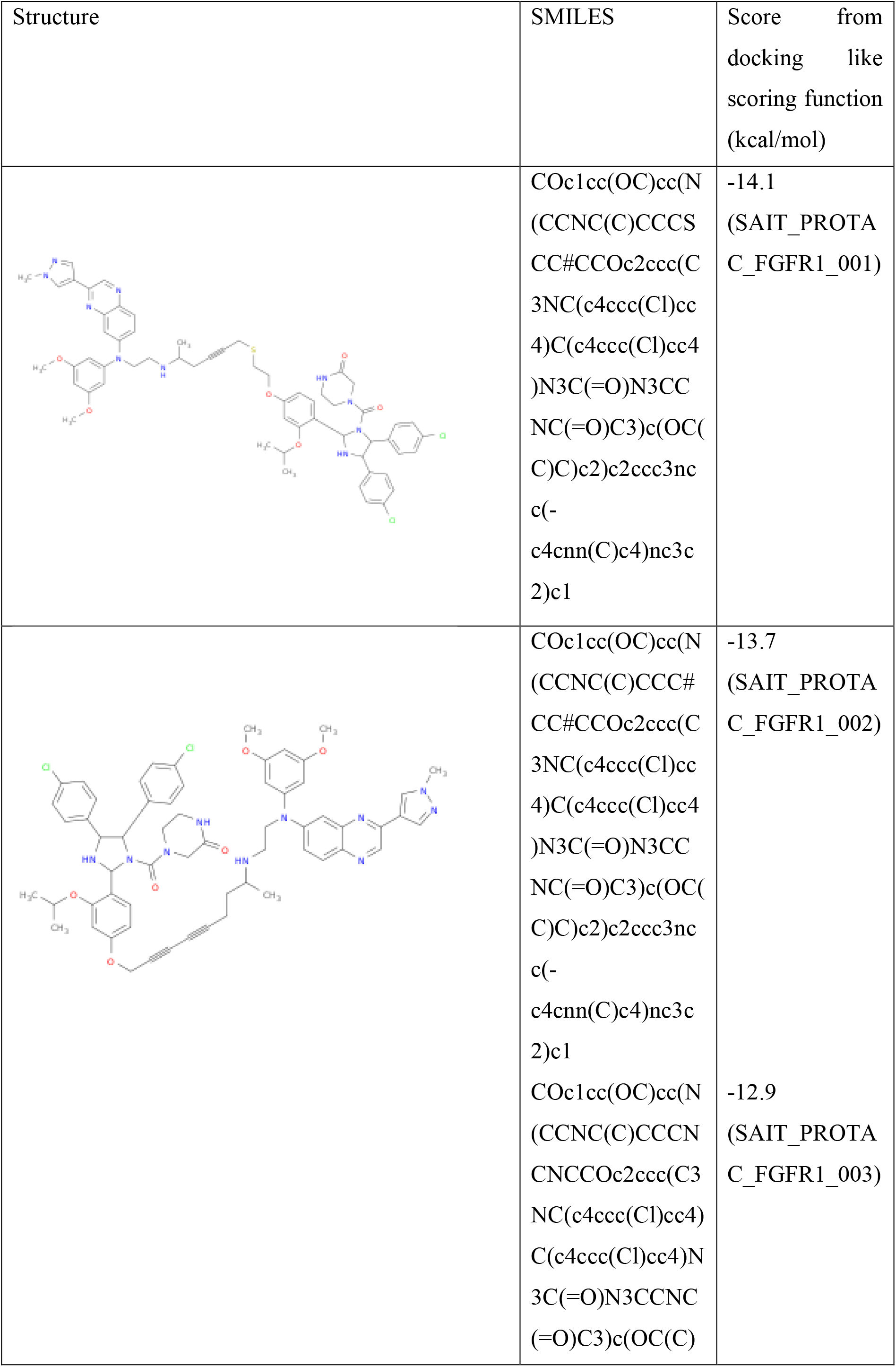
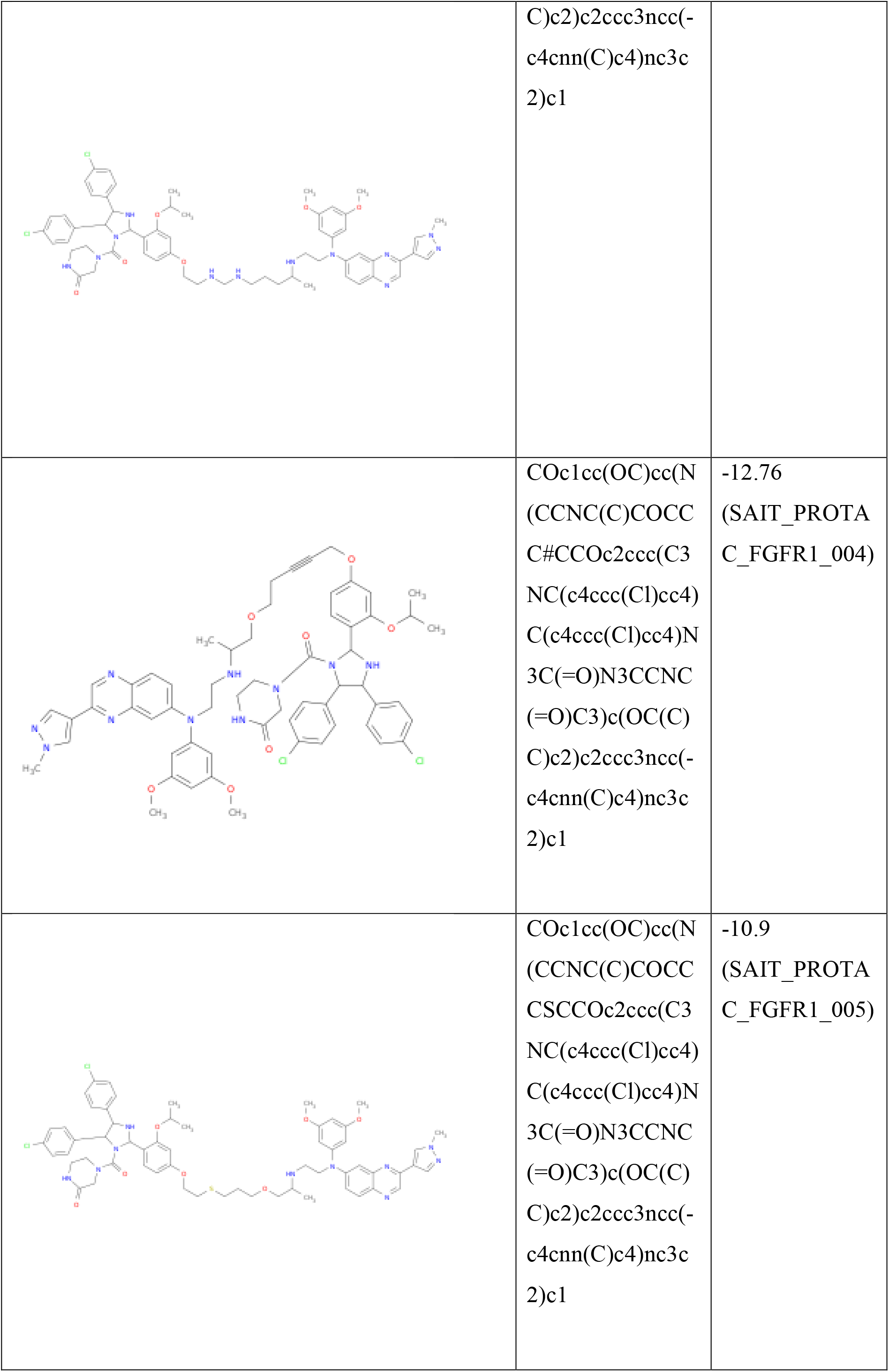
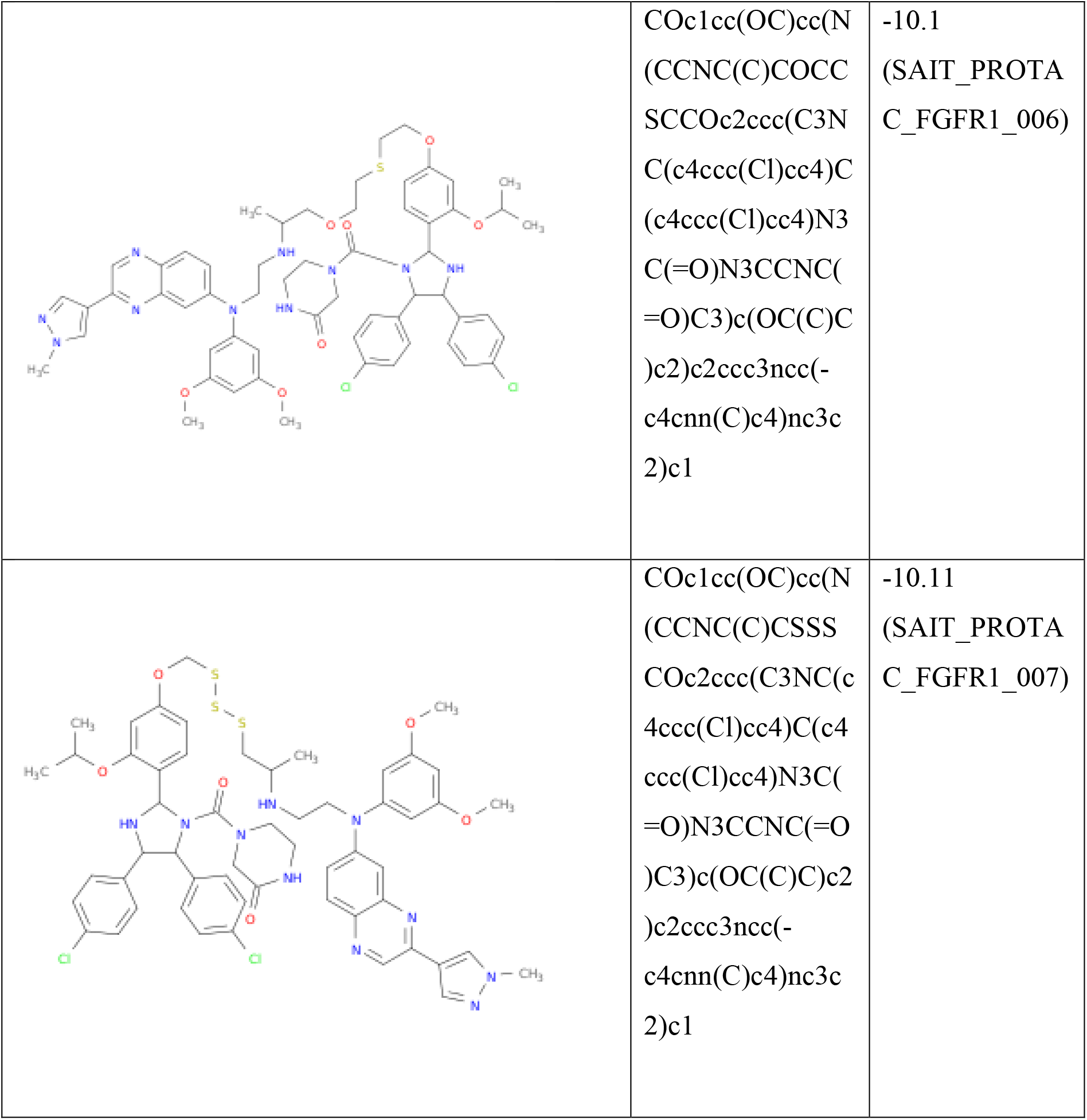
Top ranking PROTAC candidates (SAIT_PROTAC_FGFR1_001 to SAIT_PROTAC_FGFR1_007)

The top docking poses of the top ranking PROTAC candidate are shown in Fig. 3a, 3b, and 3c. The PROTAC is at the pocket which lies at the interface of FGFR-1 and MDM2. The MDM2 is shown in blue, FGFR-1 in green and the PROTAC in pink.

**Fig. 3a.**
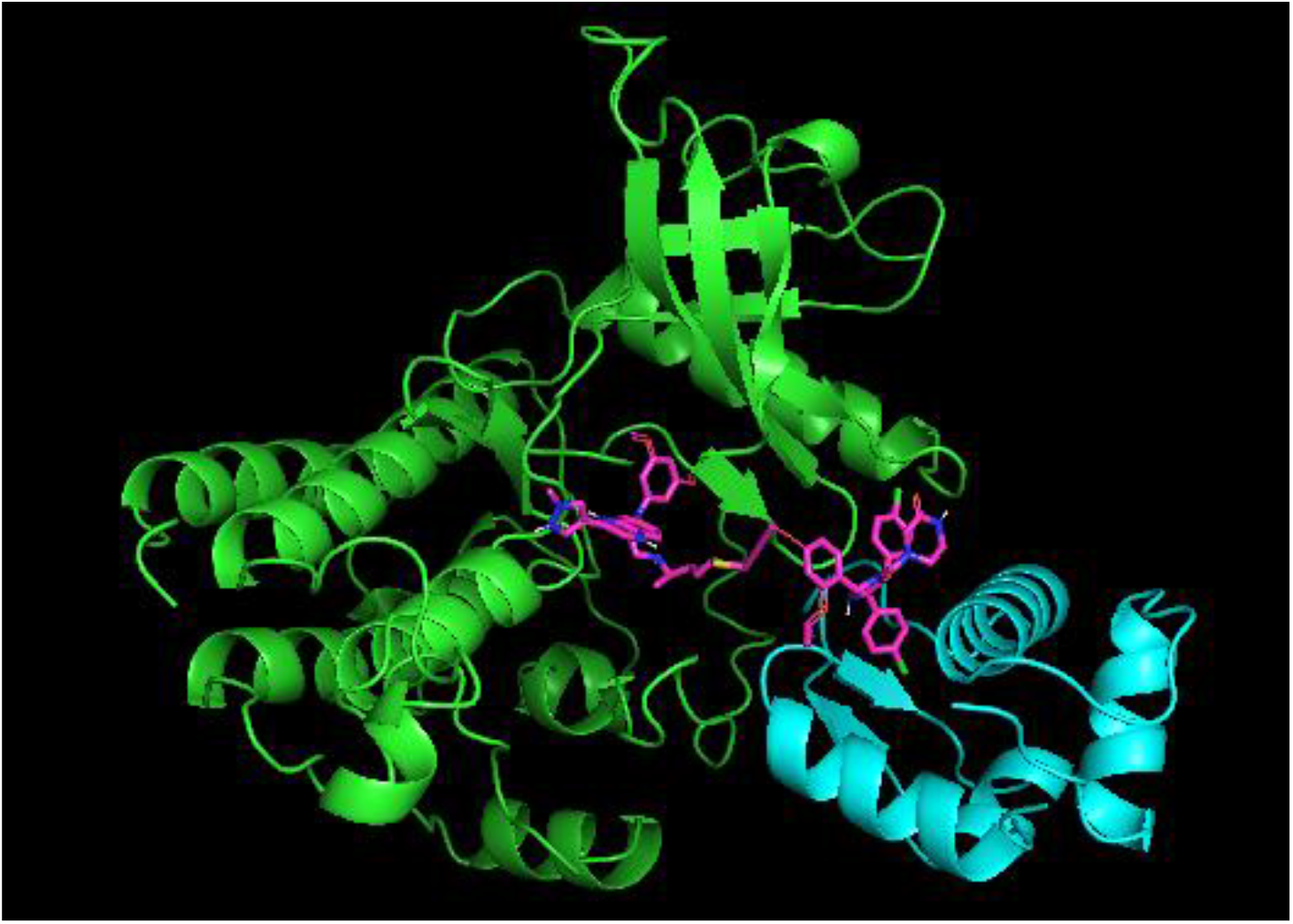
Top Pose of PROTAC in the pocket which is at the interface of FGFR1 – MDM2.

**Fig. 3b.**
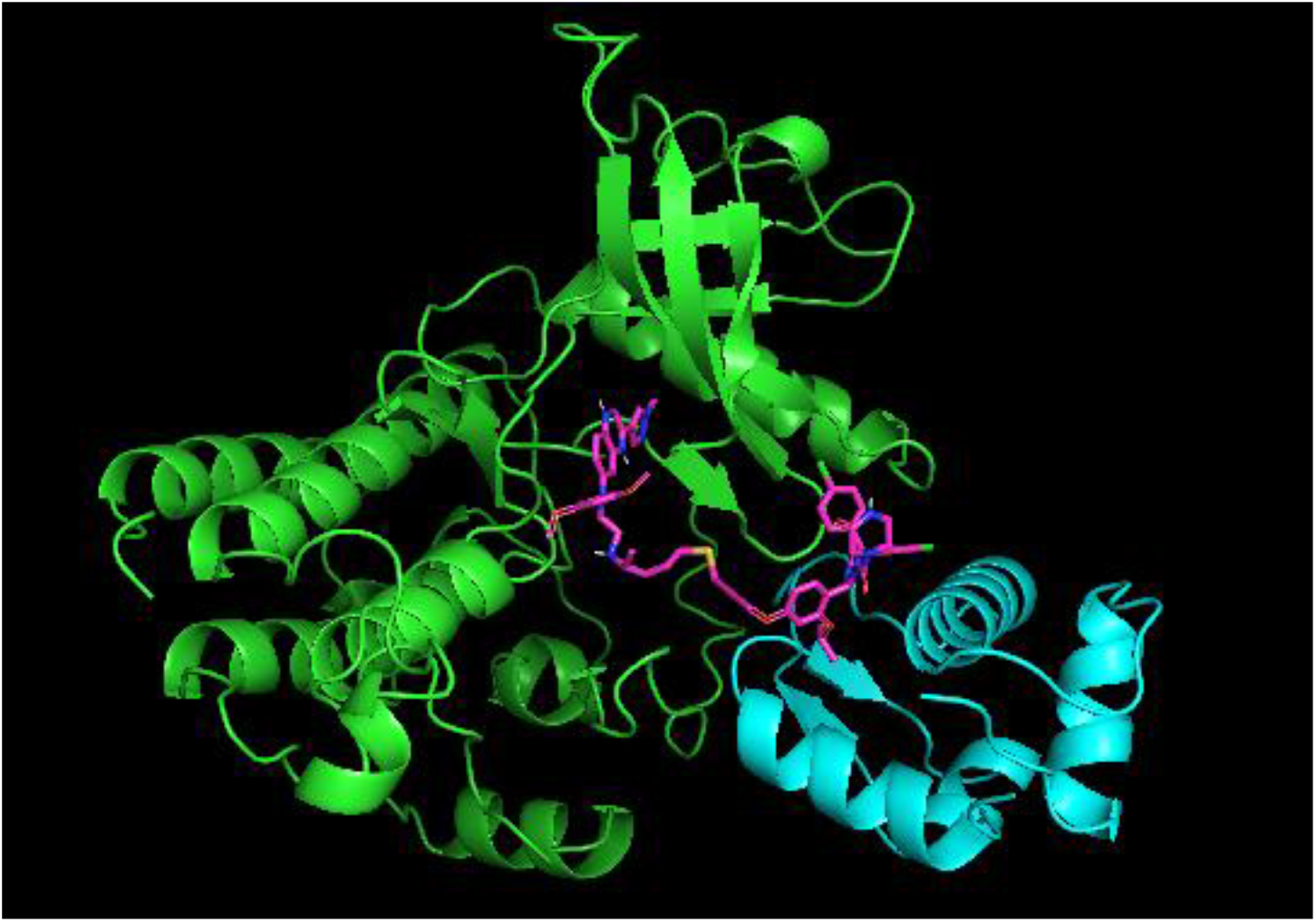
2^nd^ Pose of PROTAC in the pocket which is at the interface of FGFR1 – MDM2.

**Fig. 3c.**
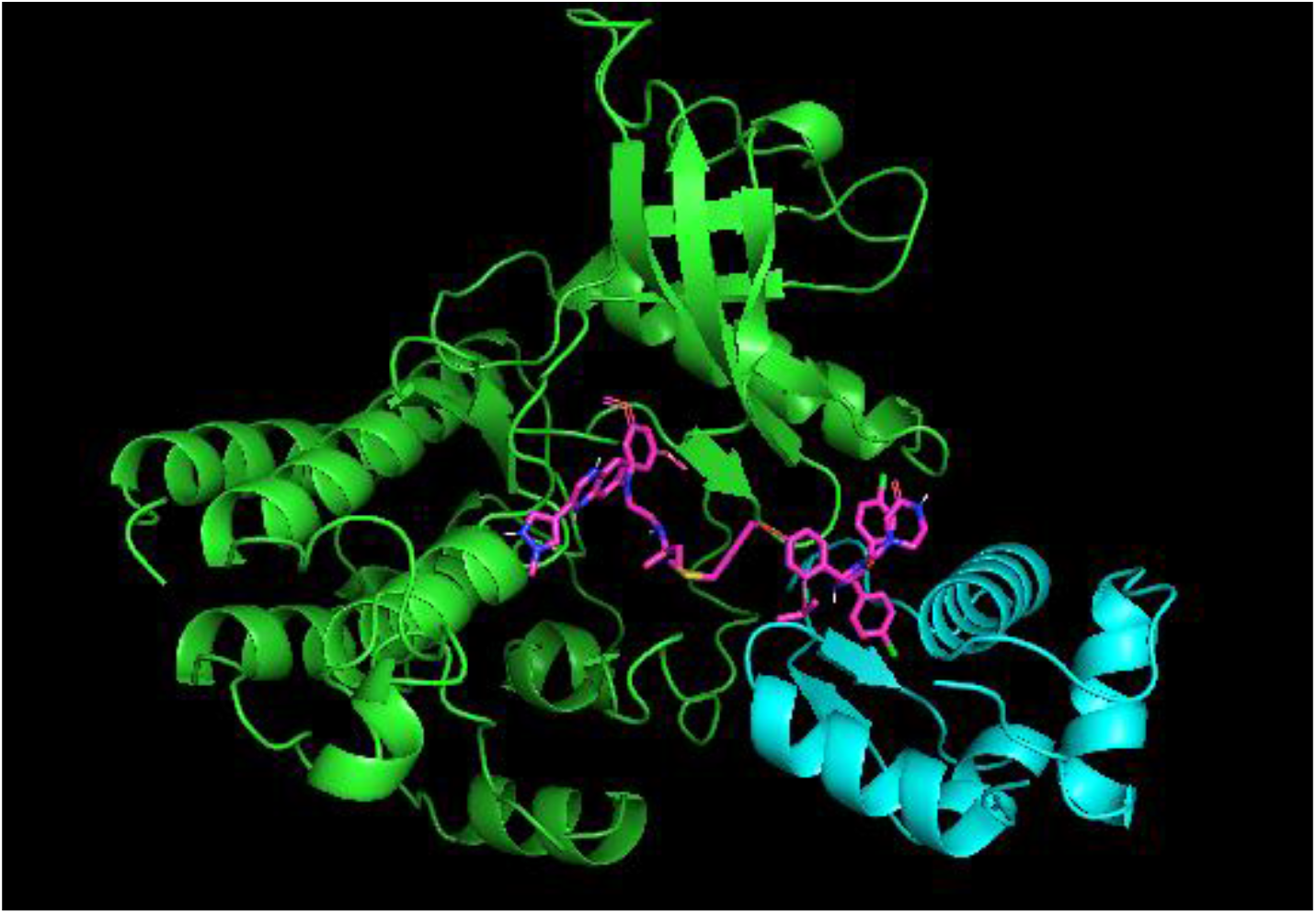
3^rd^ Pose of PROTAC in the pocket which is at the interface of FGFR1 – MDM2.

The top scoring pose (Fig. 3a) was chosen for a 200 ns long MD simulation run and it was found that the PRTOAC candidate was able to stably hold together the complex FGFR1 – MDM2. After the 200 ns long run the interactions of the PROTAC with the pocket residues of both proteins were intact, indicating stable ternary complex formation. The interacting residues from the pocket of both the proteins are shown below in Fig.4 and the interactions are listed in Table 2.

**Fig. 4.**
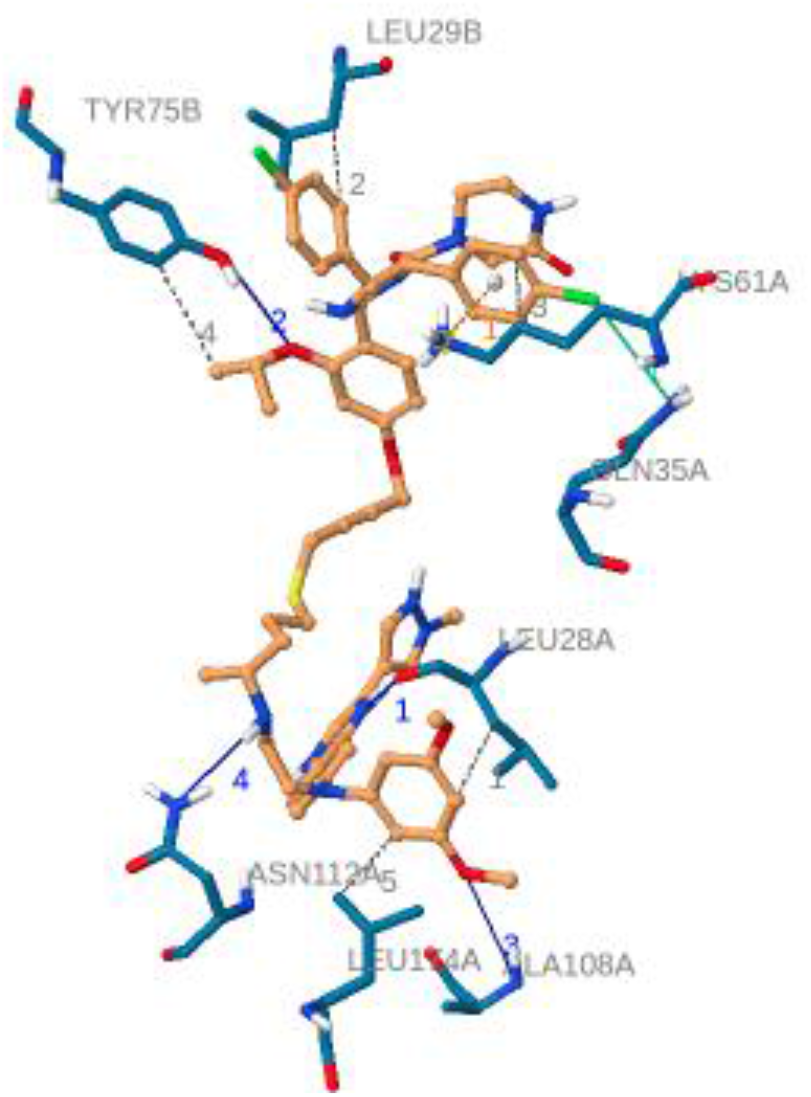
PROTAC interacting the residues from the pocket of both the proteins.

**Table 2.** PROTAC Interaction table

|  | Hydrophobic | Hydrogen bonds | Pi-Cation interaction | Halogen Bonds |
| --- | --- | --- | --- | --- |
| FGFR-1 | 28LEU, 61LYS, 174LEU | 28LEU, 108ALA, 112ASN | 61LYS | 35GLN |
| E3-ligase | 29LEU, 75TYR | 79TYR |  |  |

The results indicate that the interactions of the know binders to the proteins FGFR-1 and MDM2 are intact in the converged ternary structure after 200 ns of MD simulation.

Next, we estimated the reaction kinetics and thermodynamics using binding free energy calculations by FEP-Thermodynamic Integration method known to have very accurate results. ΔG estimated for ternary and binary complexation from FEP-TI calculations are:

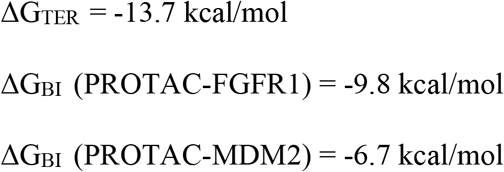

The ΔG_TER_ was found to be lower than ΔG_BI_ indicating that the ternary complexation is favoured and stable. The ΔGs we estimate theoretically are also the quantities experimentalists attempt to measure in PROTAC related assays such as ITC and SPR done to estimated ΔG for binary and ternary complexation. The FEP-TI method which is used for ΔG estimation is benchmarked in literature with a root mean square error of 0.8 kcal/mol when compared to experimental results.

## Conclusions

Through this work we contribute to a new generation of in-silico approaches to PROTAC design. To illustrate our approach, we chose FGFR-1 which is target for colorectal cancer to design a PROTAC candidate for the same. We use a generative AI based approach for PROTAC design. Among the different PROTAC candidates generated by our AI approach, we use a docking like scoring/ranking method to rank the different PROTAC candidates based on the interactions with the pocket residues of FGFR1-MDM2 and their ability to meditate the formation of the ternary complex. Further we use computationally intensive methods to estimate quantities measured in PROTAC assays which are the performance indicators for the PROTACs. The FEP-TI method which we used for ΔG estimation is benchmarked in literature with a root mean square error of 0.8 kcal/mol. The ΔG estimated through this method for ternary and binary complex indicate that the PROTAC candidate designed is able to meditate stable ternary complex formation involving FGFR1-MDM2. Based on the in-silico work conducted, we recommend synthesis and biological testing for the PROTAC candidate designed through our approach. We believe that the computational ability to calculate PROTAC performance metrics measured in PROTAC experimental assays would bring significant time reduction and cost cutting advantages to the DMTA cycle of PROTAC design. Also, in future work we see the scope of generalizing the workflow for other hetero-bifunctional therapeutics such as DUBTAC, LYTAC, AUTAC, PhosTAG (Hua et al (2022) [11]) and others where a hetero-bifunctional molecule mediated ternary complex formation is involved.

## Acknowledgements

We are grateful for the various useful discussions we have had with our colleagues at Sravathi Artificial Intelligence. In particular we would like to thank Srinivasan Krishnaswami, Raghu Bhagavat, and Nivedita Bharti for their valuable input and insights.

## References

1. Drummond, Michael L., and Christopher I. Williams. “In silico modeling of PROTAC-mediated ternary complexes: validation and application.” Journal of chemical information and modeling 59.4 (2019): 1634–1644

2. Testa, Andrea, et al. “Structure-based design of a macrocyclic PROTAC.” Angewandte Chemie 132.4 (2020): 1744–1751.

3. Liao, Junzhuo, et al. “In Silico Modeling and Scoring of PROTAC-Mediated Ternary Complex Poses.” Journal of Medicinal Chemistry 65.8 (2022): 6116–6132.

4. Li, Wenqing, et al. “Importance of Three-Body Problems and Protein–Protein Interactions in Proteolysis-Targeting Chimera Modeling: Insights from Molecular Dynamics Simulations.” Journal of Chemical Information and Modeling (2022).

5. Roy, Michael J., et al. “SPR-measured dissociation kinetics of PROTAC ternary complexes influence target degradation rate.” ACS chemical biology 14.3 (2019): 361–368.

6. Hu, Zhenyi, and Craig M. Crews. “Recent Developments in PROTAC-Mediated Protein Degradation: From Bench to Clinic.” ChemBioChem 23.2 (2022): e202100270.

7. Accurate calculation of the absolute free energy of binding for drug molecules.” Chemical science 7.1 (2016): 207–218

8. Imrie, Fergus, et al. “Deep generative models for 3D linker design.” Journal of chemical information and modeling 60.4 (2020): 1983–1995.

9. Hines, John, et al. “MDM2-recruiting PROTAC offers superior, synergistic antiproliferative activity via simultaneous degradation of BRD4 and stabilization of p53.” Cancer research 79.1 (2019): 251–262

10. Rezaei, Mohammad Ali, Yanjun Li, Dapeng Wu, Xiaolin Li, and Chenglong Li. “Deep learning in drug design: protein-ligand binding affinity prediction.” IEEE/ACM Transactions on Computational Biology and Bioinformatics (2020).

11. Hua, Liwen, Qiuyue Zhang, Xinyue Zhu, Ruoning Wang, Qidong You, and Lei Wang. “Beyond Proteolysis-Targeting Chimeric Molecules: Designing Heterobifunctional Molecules Based on Functional Effectors.” Journal of Medicinal Chemistry (2022).

