## Supplementary Information for "A New In-Silico Approach for PROTAC Design and Quantitative Rationalization of PROTAC mediated Ternary Complex Formation"

**SUPLEMENTARY INFORMATION**

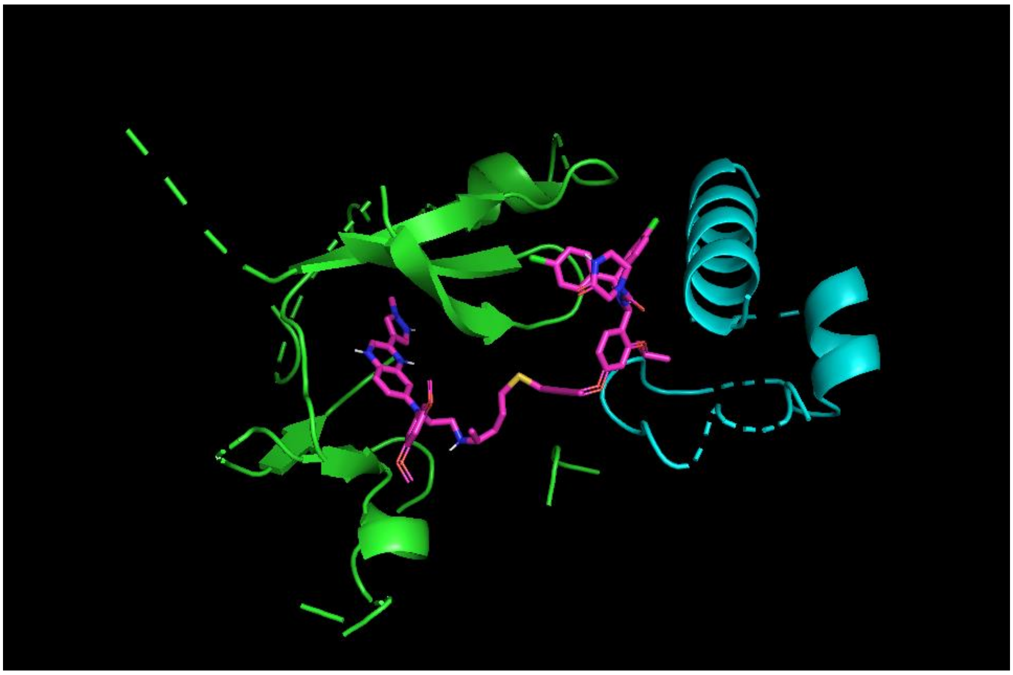

Fig.1 – Pocket residues of FGFR1-MDM2 and a PROTAC conformer

**Protein-protein docking of FGFR1 – MDM2 and selecting a low energy docked pose of FGFR1 – MDM2 wherein the known binders in their respective pockets can be connected by a linker**

↓

**Deep Generative linker design and generation of multiple PROTAC candidates**

↓

**Docking based scoring and ranking of PROTAC candidates**

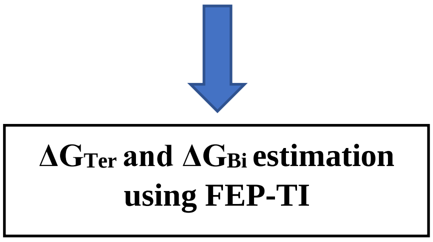


Fig.2 Workflow diagram.

Table 1 – Top ranking PROTAC candidates (SAIT_PROTAC_FGFR1_001 to SAIT_PROTAC_FGFR1_007)

| Structure | SMILES | Score from docking like scoring function (kcal/mol) |
|---|---|---|
| 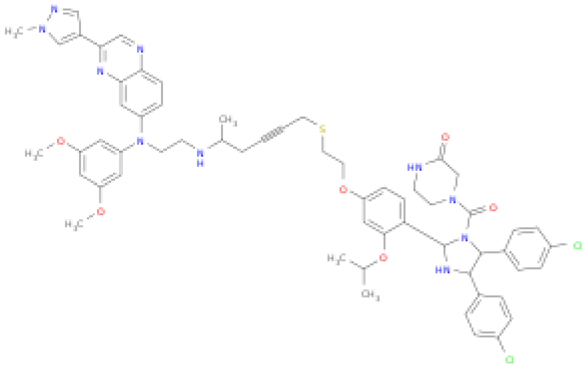 | COc1cc(OC)cc(N(CCNC(C)CCCSCC#CCOc2ccc(C3NC(c4ccc(Cl)cc4)C(c4ccc(Cl)cc4)N3C(=O)N3CCNC(=O)C3)c(OC(C)C)c2)c2ccc3ncc(-c4cnn(C)c4)nc3c2)c1 | -14.1 (SAIT_PROTAC_FGFR1_001) |

| | | |
|---|---|---|
| 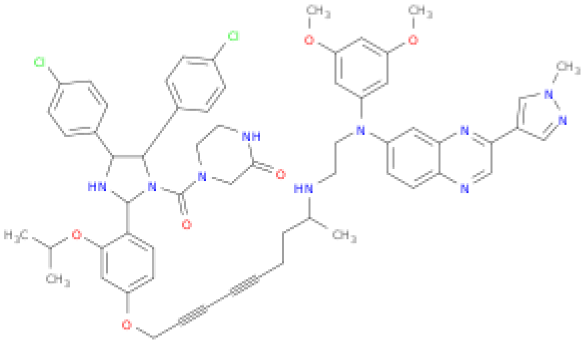 | COc1cc(OC)cc(N(CCNC(C)CCC#CC#CCOc2ccc(C3NC(c4ccc(Cl)cc4)C(c4ccc(Cl)cc4)N3C(=O)N3CCNC(=O)C3)c(OC(C)C)c2)c2ccc3ncc(-c4cnn(C)c4)nc3c2)c1 | -13.7 (SAIT_PROTAC_FGFR1_002) |
| 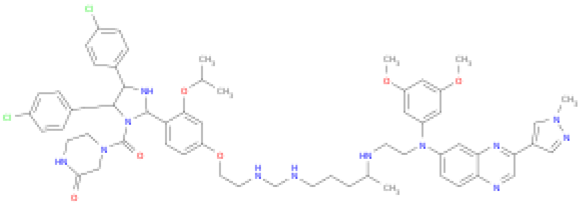 | COc1cc(OC)cc(N(CCNC(C)CCCNCNCCOc2ccc(C3NC(c4ccc(Cl)cc4)C(c4ccc(Cl)cc4)N3C(=O)N3CCNC(=O)C3)c(OC(C)C)c2)c2ccc3ncc(-c4cnn(C)c4)nc3c2)c1 | -12.9 (SAIT_PROTAC_FGFR1_003) |

| | | |
|---|---|---|
| 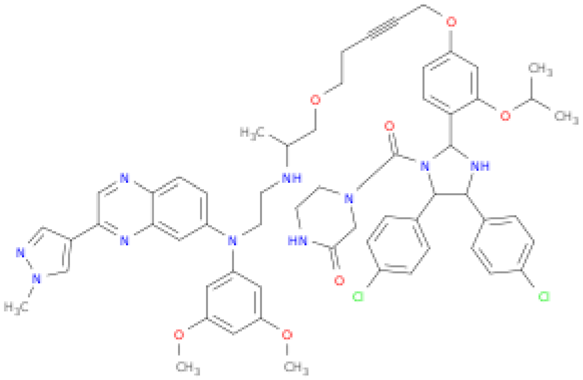 | COc1cc(OC)cc(N(CCNC(C)COCCC#CCOc2ccc(C3NC(c4ccc(Cl)cc4)C(c4ccc(Cl)cc4)N3C(=O)N3CCNC(=O)C3)c(OC(C)C)c2)c2ccc3ncc(-c4cnn(C)c4)nc3c2)c1 | -12.76 (SAIT_PROTAC_FGFR1_004) |
| 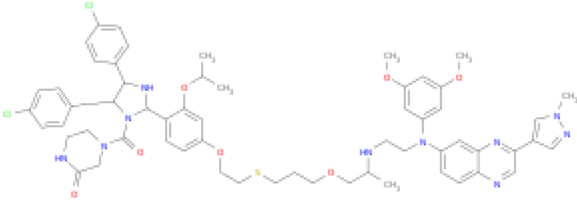 | COc1cc(OC)cc(N(CCNC(C)COCCCSCCOc2ccc(C3NC(c4ccc(Cl)cc4)C(c4ccc(Cl)cc4)N3C(=O)N3CCNC(=O)C3)c(OC(C)C)c2)c2ccc3ncc(-c4cnn(C)c4)nc3c2)c1 | -10.9 (SAIT_PROTAC_FGFR1_005) |
| 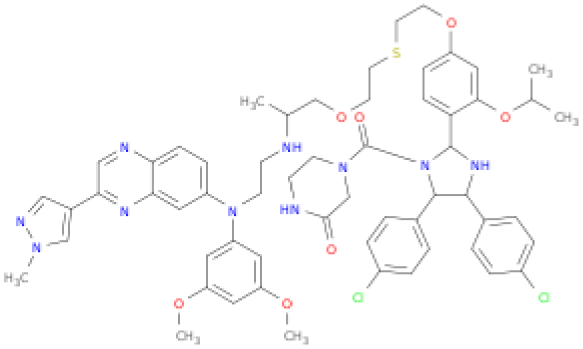 | COc1cc(OC)cc(N(CCNC(C)COCCSCCOc2ccc(C3NC(c4ccc(Cl)cc4)C(c4ccc(Cl)cc4)N3C(=O)N3CCNC(=O)C3)c(OC(C)C)c2)c2ccc3ncc(-c4cnn(C)c4)nc3c2)c1 | -10.1 (SAIT_PROTAC_FGFR1_006) |

| | | |
|---|---|---|
| 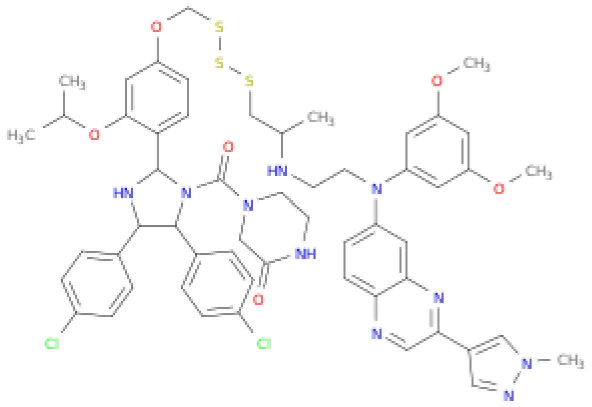 | COc1cc(OC)cc(N(CCNC(C)CSSSCOc2ccc(C3NC(c4ccc(Cl)cc4)C(c4ccc(Cl)cc4)N3C(=O)N3CCNC(=O)C3)c(OC(C)C)c2)c2ccc3ncc(-c4cnn(C)c4)nc3c2)c1 | -10.11 (SAIT_PROTAC_FGFR1_007) |

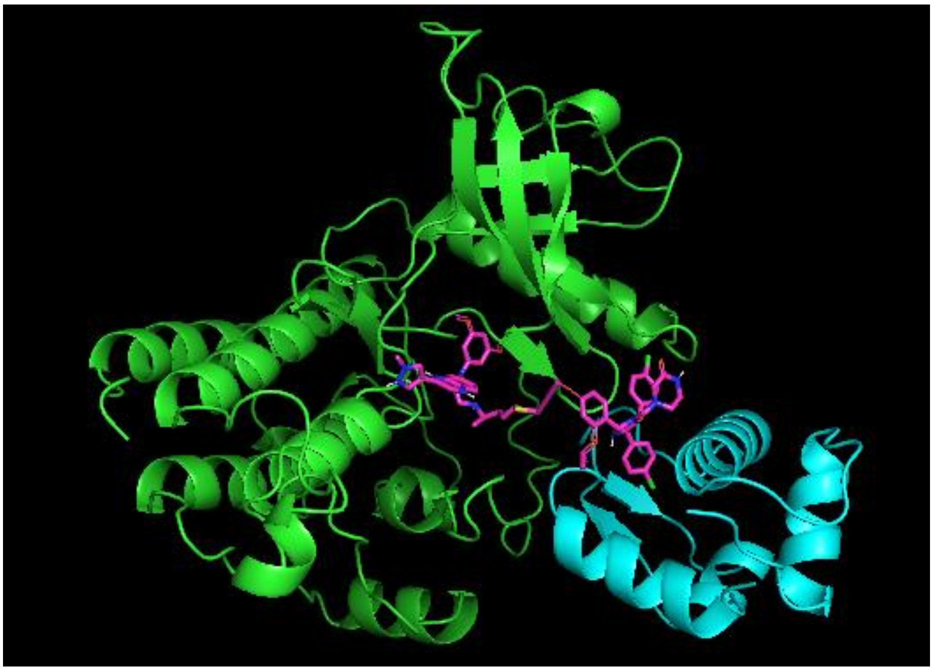

Fig.3a – Top Pose of PROTAC in the pocket which is at the interface of FGFR1 – MDM2.

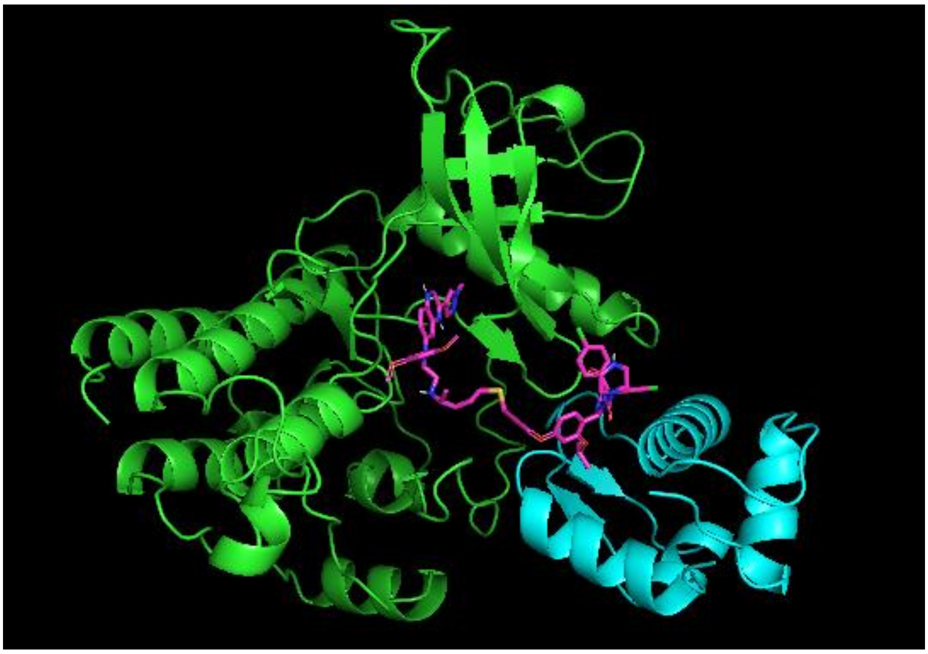

Fig.3b– 2nd Pose of PROTAC in the pocket which is at the interface of FGFR1 – MDM2.

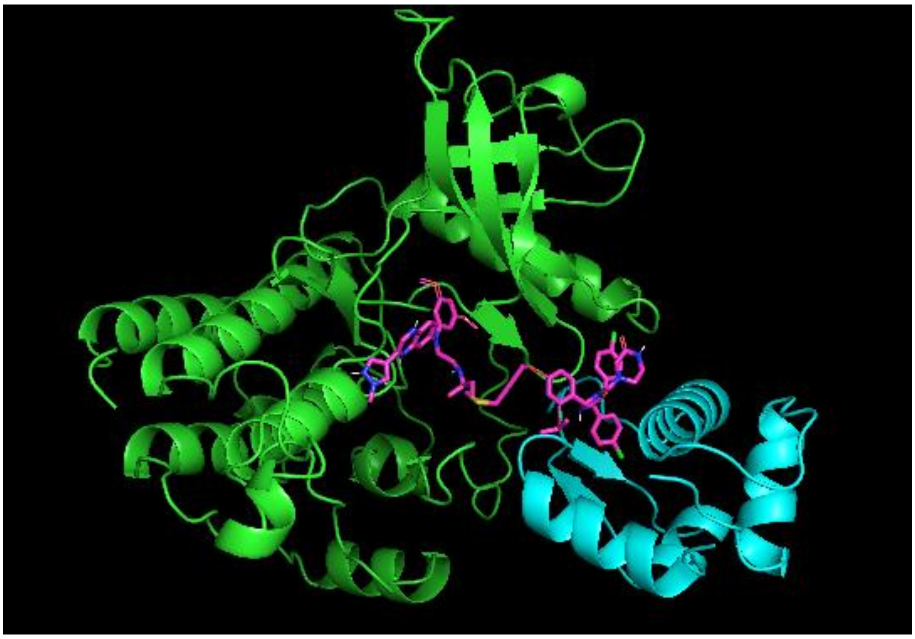

Fig.3c – 3rd Pose of PROTAC in the pocket which is at the interface of FGFR1 – MDM2.

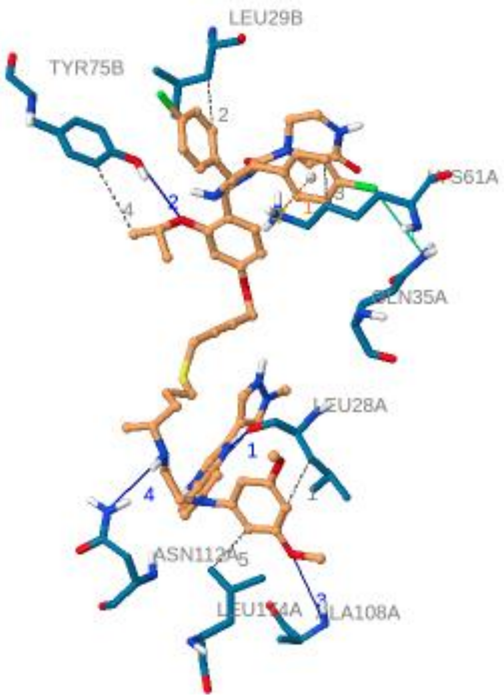


Fig. 4 – PROTAC interacting the residues from the pocket of both the proteins.

Table 2 – PROTAC Interaction table

| | Hydrophobic | Hydrogen bonds | Pi-Cation interaction | Halogen Bonds |
|---|---|---|---|---|
| FGFR-1 | 28LEU, 61LYS, 174LEU | 28LEU, 108ALA, 112ASN | 61LYS | 35GLN |
| E3-ligase | 29LEU, 75TYR | 79TYR | | |
